# Distinct cerebrospinal fluid DNA methylation signatures linked to Alzheimer’s disease

**DOI:** 10.1101/2025.04.03.647115

**Authors:** Magdalena Rados, Sigrid Klotz, Günther Regelsberger, Elisabeth Stögmann, Lukas D. Landegger, Irfete S. Fetahu

## Abstract

Alzheimer’s disease (AD) accounts for more than 60% of the dementia cases and currently there is no curative treatment for it. With the emergence of potentially disease modifying treatments, early diagnosis is key to identify patient groups that would benefit from such treatments, aiming to prevent severe cognitive decline. We previously identified a set of DNA methylation signatures that allow for accurate diagnosis of AD in cortical neurons and brain tissue, even before clinical manifestation of the disease [1]. Here we investigate 11 of these signature regions via targeted next-generation sequencing in cell-free DNA (cfDNA) isolated from cerebrospinal fluid (CSF) of AD patients homozygous for *APOE4* (n=4) and sporadic AD (n=5) cases compared to age-matched control samples (n=5). Our analyses demonstrated that 6/11 of the tested DNA methylation signatures that had initially been identified in cortical neurons and brain tissue were also validated in cfDNA. The remainder of the tested regions either showed opposite trends (3/11) or did not result in any differences (2/11) between control and AD cases. Thus, this presents a direct approach allowing to test for these DNA methylation signatures in CSF-derived cfDNA, and bypasses the need to generate induced pluripotent stem cell-derived cortical neurons from patients.

## Main

AD is a complex and multifactorial neurodegenerative disease characterized by a long prodromal phase. However, early diagnosis of AD is pivotal in disease management given that currently only symptomatic treatments exist [2]. The genetic factors currently linked with AD provide a platform for screening for familial AD [3], but most AD cases do not have a well-defined etiology. Detection of biomarkers, such as amyloid-beta and total (t) or phosphorylated (p) Tau proteins in CSF allows for an accurate diagnosis in tandem with clinical evaluation.

However, during prodromal stages only changes in amyloid-beta are noted, but this is not sufficient to discern AD from other types of dementia. Presence of tTau and pTau proteins has been shown to aid in differential diagnosis between AD and other types of dementia [4], but pTau levels increase only in advanced stages of the disease [5]. Therefore, finding biomarkers that apply to all AD pathogenesis still remains a diagnosis dilemma, where early and accurate characterization is vital to enable a timely intervention, potentially slowing the progression of AD.

Altered epigenome, in particular aberrant methylation of DNA CpG sites is virtually present in any disease. DNA methylation is critical in neuronal development and maturation, and several studies have reported aberrant DNA methylation in a host of neural disorders, including AD [6, 7]. We have previously reported a set of DNA methylation signatures that can accurately diagnose all AD cases (98% specificity), long before clinical manifestation of the disease, and that these signatures are age-independent [1]. Here, we apply those signatures in cfDNA from CSF of controls and AD patients, where we recapitulate the majority of our previous findings, circumventing the need for brain tissue or the use of patient stem-cell-derived neurons.

DNA methylation at the 5th position on cytosine (5mC) is the most abundant form of DNA methylation, therefore, here we focused only on the 5mC signatures. To test this, we designed a targeted DNA methylation PCR panel, covering CpG sites that had been identified in our previous work (**Supplementary Table 1**) [1]. Following assay validation, we measured the 5mC levels in our cohort samples, which included a total of nine AD patients that were selected based on biomarkers, including amyloid-beta 42, tTau, and pTau proteins (**Fig. 1** and **Table 1**) coupled with clinical symptoms (**Supplementary Table 2**). Patient characteristics and clinical symptoms were collected retrospectively from clinical files. As controls, we included three age-matched samples that were from patients with normal pressure hydrocephalus with normal CSF AD biomarker profiles. We also included two samples from vestibular schwannoma patients, a benign tumor arising from Schwann cells surrounding the vestibulocochlear nerve, where neurodegeneration is generally uncommon [8]. Our data showed that DNA methylation levels in CSF-derived cfDNA of the following genes: *KIF26A, NFIX, PCDHA2, PTPRN2* (previously annotated as miR-153) were marked by loss of methylation (**Fig. 2a**) whereas *NACC2* and *NFATC1* were characterized by gain of methylation (**Fig. 2b**). These data recapitulate the trends initially identified in brain tissues and cortical neurons generated from patient-derived stem cells [1]. The rest of the tested genes either followed opposite trends, *ADA2, CASZ1, LINC02055* (**Supplementary Fig.1a**) or did not show any variations between control and AD samples, *NR4A2, PKHD1* (**Supplementary Fig.1b**) compared to the previous data employing cortical neurons and brain tissue [1]. Finally, genes *KIF26A, PTPRN2, NACC2, NFATC1* validated herein in CSF samples were critical to accurately and consistently (98% specificity, 48% sensitivity) delineate between control and AD tissue samples in our DNA methylation data [1], and in the DNA methylation data from the Religious Orders Study/Memory and Aging Project [9], the largest longitudinal AD study to date [10]. Therefore, we propose the use of DNA methylation levels of these six genes and their further validation as AD biomarkers that can be tested in CSF samples in patients suspected of having AD, for early treatment intervention, ultimately aiming to mitigate severe cognitive decline, and hopefully provide a platform for new treatment modalities. Finally, it would be relevant to investigate whether these epigenetic markers are limited to AD or extend to other neurodegenerative diseases.

**Table 1.**
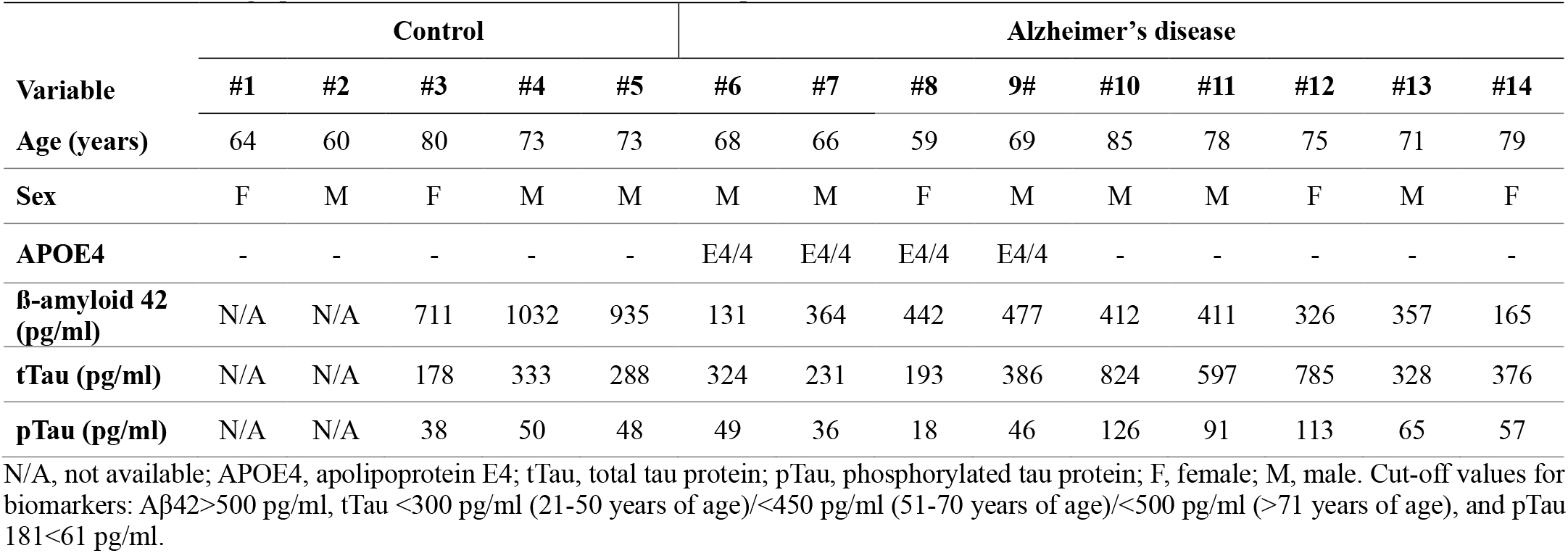
Cohort demographics of controls and Alzheimer’s disease patients.

| Variable | Control |  |  |  |  | Alzheimer's disease |  |  |  |  |  |  |  |  |
| --- | --- | --- | --- | --- | --- | --- | --- | --- | --- | --- | --- | --- | --- | --- |
|  | #1 | #2 | #3 | #4 | #5 | #6 | #7 | #8 | 9# | #10 | #11 | #12 | #13 | #14 |
| Age (years) | 64 | 60 | 80 | 73 | 73 | 68 | 66 | 59 | 69 | 85 | 78 | 75 | 71 | 79 |
| Sex | F | M | F | M | M | M | M | F | M | M | M | F | M | F |
| APOE4 | - | - | - | - | - | E4/4 | E4/4 | E4/4 | E4/4 | - | - | - | - | - |
| β-amyloid 42 (pg/ml) | N/A | N/A | 711 | 1032 | 935 | 131 | 364 | 442 | 477 | 412 | 411 | 326 | 357 | 165 |
| tTau (pg/ml) | N/A | N/A | 178 | 333 | 288 | 324 | 231 | 193 | 386 | 824 | 597 | 785 | 328 | 376 |
| pTau (pg/ml) | N/A | N/A | 38 | 50 | 48 | 49 | 36 | 18 | 46 | 126 | 91 | 113 | 65 | 57 |

N/A, not available; APOE4, apolipoprotein E4; tTau, total tau protein; pTau, phosphorylated tau protein; F, female; M, male. Cut-off values for biomarkers: Aβ42>500 pg/ml, tTau <300 pg/ml (21-50 years of age)/<450 pg/ml (51-70 years of age)/<500 pg/ml (>71 years of age), and pTau 181<61 pg/ml.

**Fig. 1.**
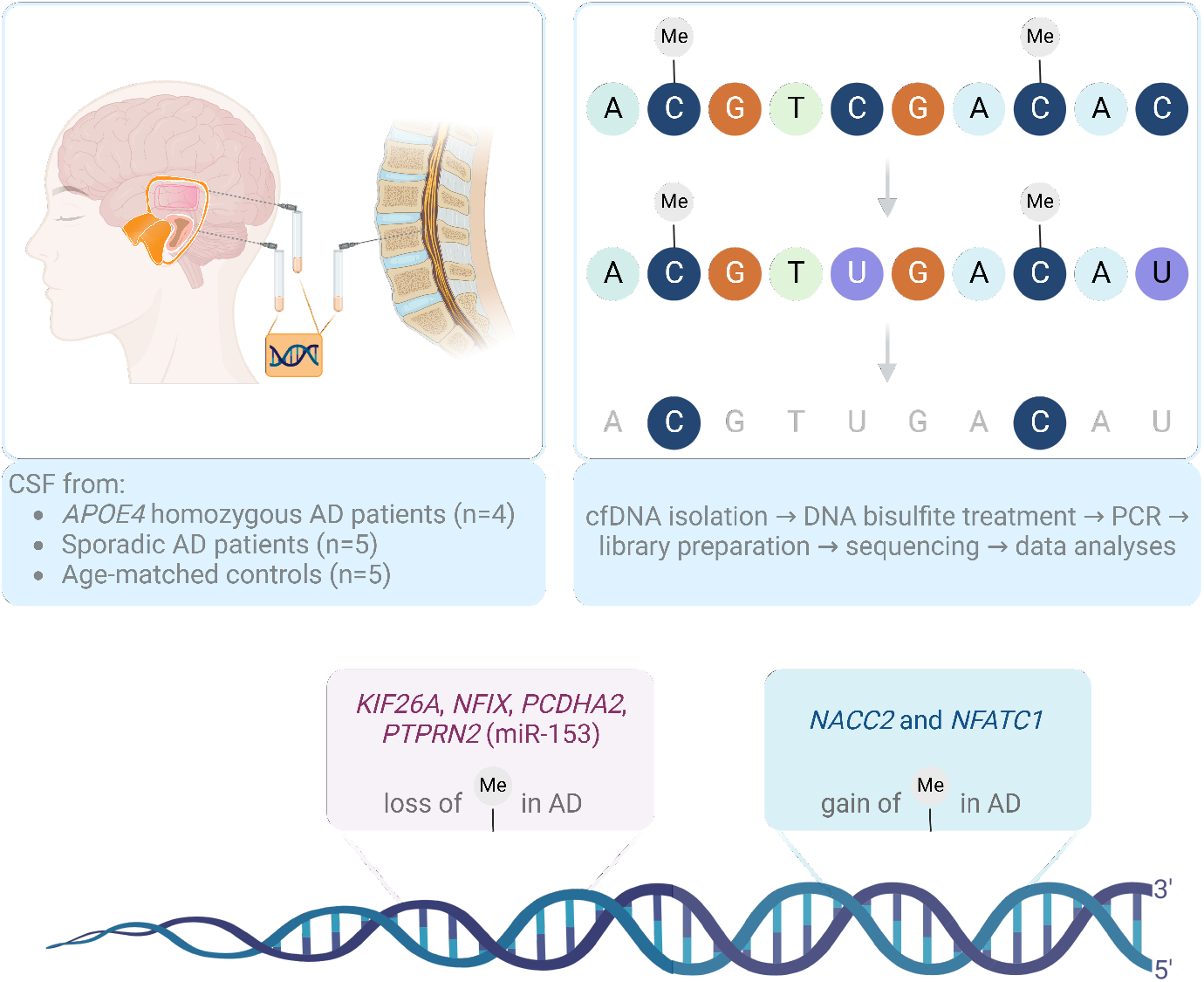
Schematic illustration of the testing for DNA methylation signatures in cfDNA isolated from CSF. The figure depicts the methodological approach and summarizes the main findings. AD, Alzheimer’s disease; APOE4, apolipoprotein E 4; cfDNA, cell-free DNA; CSF, cerebrospinal fluid.

**Fig. 2.**
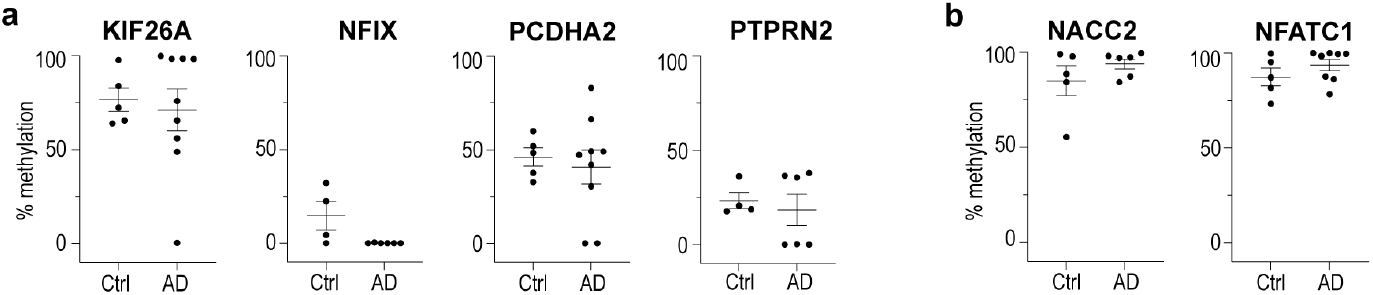
DNA methylation levels of gene signatures associated with Alzheimer’s disease tested in CSF. Targeted bisulfite sequencing of genes linked to loss (a) and gain (b) of methylation in AD patients (n=6-9 biologically independent samples) compared to controls (n=4-5 biologically independent samples). The data are presented as the mean of methylation levels, if more than one CpG/gene site was interrogated. Medians are indicated by the lines.

## Material and methods

### Collection of cerebrospinal fluid samples

All patient material (**Table 1** and **Supplementary Table 2**) used in this study was obtained from the Vienna General Hospital after written informed consent was obtained from patients for the use of left-over samples for research. Ethical approval for the Neurology/Neuropathology and Neurochemistry, and Neuro-Oncology Biobanks as well as use of clinical data was obtained from the local institutional review board of the Medical University of Vienna (EK: 1636/2019, EK: 1454/2018, EK: 1375/2018). The CSF was either obtained via lumbar puncture as per standard practice or intraoperatively either via the middle fossa or the translabyrinthine approaches during vestibular schwannoma resection. Samples were collected in polypropylene tubes and then were centrifuged at 4,200g for 3 minutes, following which supernatant was collected, aliquoted, and stored at –80°C until further use. Due to limited sample size of biological materials collected from patients, these materials are not available to be shared.

### Quantification of B-amyloid, total tau, and phosphorylated tau

The levels of Aβ42 [INNOTEST β-AMYLOID (1-42)], tTau (INNOTEST hTAU-Ag), and pTau 181 [INNOTEST PHOSPHO-TAU (181P)] were determined in undiluted CSF samples by ELISA, all from Fujirebio, the FDA approved kit according to the manufacturer’s instructions. Samples were processed in duplicates and absorbance was measured at 540nm (reference wavelength 620 nm) within 15 min after addition of the stop solution using PowerWave microplate spectrophotometer XS2 and the Gen5 software (BioTek). Standard curves were generated in parallel for each assay. The cut-off values for the biomarkers were based on the manufacturer’s recommendation: Aβ42>500 pg/ml, tTau <300 pg/ml (21-50 years of age)/<450 pg/ml (51-70 years of age)/<500 pg/ml (>71 years of age), and pTau 181<61 pg/ml.

### DNA isolation and library preparation

DNA isolation was performed using 500μL of CSF employing QIAamp DNA Mini Kit (Qiagen) following the manufacturer’s instructions. DNA samples were bisulfite treated using the EZ-96 DNA methylation kit (Zymo Research) and subjected to simplex or multiplex PCRs using 0.5 units of HotStarTaq DNA polymerase (Qiagen), 0.2μM primers, and 3μL of bisulfite-treated DNA in a 20μL reaction. Prior to library preparation, PCR products from the same sample were pooled and then purified using the QIAquick PCR Purification Kit columns or plates (Qiagen). All PCR products were verified using the Qiagen QIAxcel Advanced System (v1.0.6). Target samples were run alongside established reference DNA samples with a range of methylation (0, 5, 10, 25, 50, 75, and 100% methylation) and library generation was performed by EpigenDx, MA, USA. Next, library molecules were purified using Agencourt AMPure XP beads (Beckman Coulter). Barcoded samples were then pooled in an equimolar fashion before template preparation and enrichment were performed on the Ion Chef™ system using Ion 520™ & Ion 530™ ExT Chef reagents (Thermo Fisher). Following this, enriched, template-positive library molecules were sequenced on the Ion S5™ sequencer using an Ion 530™ sequencing chip.

## Data analyses

FASTQ files from the Ion Torrent S5 server were aligned to a local reference database using the open-source Bismark Bisulfite Read Mapper program (v0.12.2) with the Bowtie2 alignment algorithm (v2.2.3). Methylation levels were calculated in Bismark by dividing the number of methylated reads by the total number of reads. An R-squared value (RSQ) was calculated from the controls set at known methylation levels to test for PCR bias.

## Acknowledgements

This study was supported by the Austrian Science Fund (FWF#P35072) to I.S.F and the Departments of Neurology and Otolaryngology of the Medical University of Vienna. We thank all the patients who participated in this study. We are grateful to Drs. Ellen Gelpi and Romana Höftberger for the fruitful scientific discussions.

## Contributions

I.S.F., conceived, designed, and supervised the execution of the entire study. M.R. and I.S.F. performed experiments pertinent to the DNA methylation studies, analyzed, and interpreted the data. G.R., conducted the ELISA experiments and analyzed the data. S.K., E.S., and L.D.L contributed samples, clinical data, and annotated the cohort. I.S.F. wrote the manuscript with contributions from all authors. All authors have revised the manuscript, have read and agreed to the published version of the manuscript.

---

**Supplementary Fig. 1.**
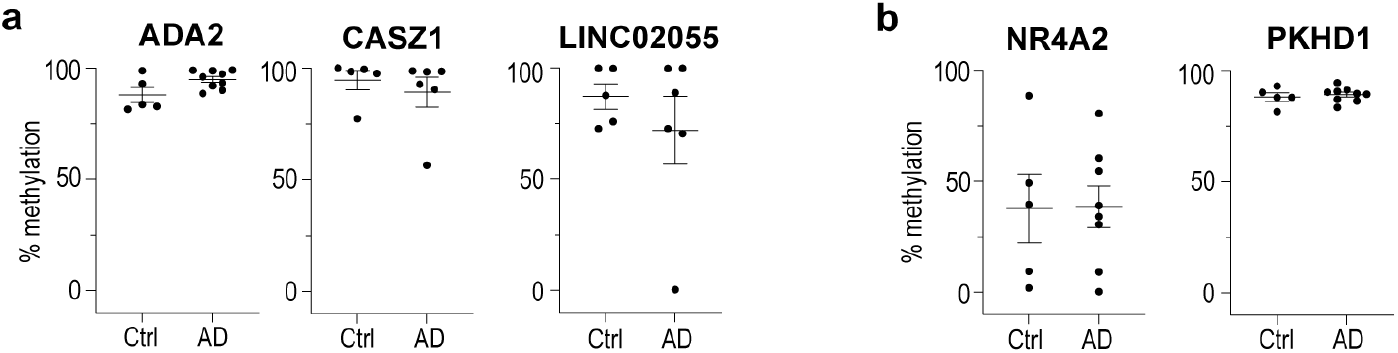
DNA methylation levels of Alzheimer’s disease gene signatures tested in CSF. (a) Targeted bisulfite sequencing of genes following opposite changes (compared to the original study) as shown in (a) and (b) no differences in methylation levels in AD patients (n=6-9 biologically independent samples) compared to controls (n=4-5 biologically independent samples). The data are presented as the mean of methylation levels, if more than one CpG/gene site was interrogated. Medians are indicated by the lines.

**Supplementary Table 1.**
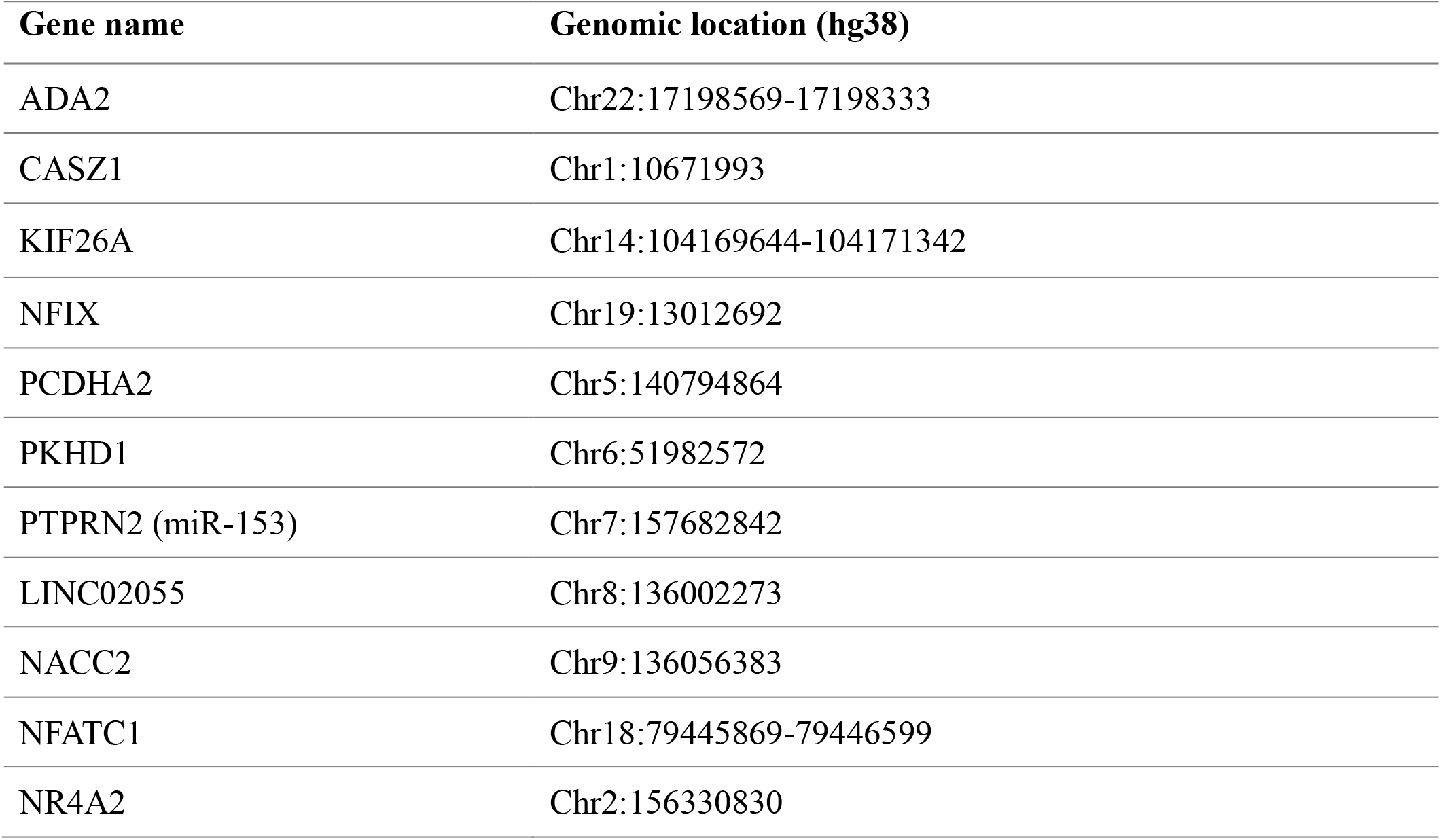
5mC methylation signatures in Alzheimer’s disease and location of CpG sites.

| **Gene name** | **Genomic location (hg38)** |
| --- | --- |
| ADA2 | Chr22:17198569-17198333 |
| CASZ1 | Chr1:10671993 |
| KIF26A | Chr14:104169644-104171342 |
| NFIX | Chr19:13012692 |
| PCDHA2 | Chr5:140794864 |
| PKHD1 | Chr6:51982572 |
| PTPRN2 (miR-153) | Chr7:157682842 |
| LINC02055 | Chr8:136002273 |
| NACC2 | Chr9:136056383 |
| NFATC1 | Chr18:79445869-79446599 |
| NR4A2 | Chr2:156330830 |

**Supplementary Table 2.** Clinical data of controls and Alzheimer’s disease patients.

| **Case** | **Clinical Symptoms** | **Group** |
| --- | --- | --- |
| 1 | Vestibular schwannoma | control |
| 2 | Vestibular schwannoma | control |
| 3 | Normal pressure hydrocephalus | control |
| 4 | Normal pressure hydrocephalus | control |
| 5 | Gait disturbance, memory problems, suspicion of normal pressure hydrocephalus | control |
| 6 | Dementia | ApoE4/4 |
| 7 | Suspicion of Alzheimer's disease, Internal hydrocephalus, cognitive decline | ApoE4/4 |
| 8 | Suspicion of Alzheimer's disease, presenile dementia, brain atrophy | ApoE4/4 |
| 9 | Dementia of unknown cause | ApoE4/4 |
| 10 | Suspicion of Alzheimer's Disease | sporadic AD |
| 11 | Dementia | sporadic AD |
| 12 | Memory problems, personality change | sporadic AD |
| 13 | Rapidly progressive dementia | sporadic AD |
| 14 | Personality change | sporadic AD |

